# Artificial neural networks for monitoring network optimisation—a practical example using a national insect survey

**DOI:** 10.1101/2020.05.14.095539

**Authors:** Yoann Bourhis, James R. Bell, Frank van den Bosch, Alice E. Milne

## Abstract

Monitoring networks are improved by additional sensors. Optimal configurations of sensors give better representations of the process of interest, maximising its exploration while minimising the need for costly infrastructure. By modelling the monitored process, we can identify gaps in its representation, *i.e.* uncertain predictions, where additional sensors should be located. Here, with data collected from the Rothamsted Insect Survey network, we train an artificial neural network to predict the seasonal aphid arrival from environmental variables. We focus on estimating prediction uncertainty across the UK to guide the addition of a sensor to the network. We first illustrate how to estimate uncertainty in neural networks, hence making them relevant for model-based monitoring network optimisation. Then we highlight critical areas of agricultural importance where additional traps would improve decision support and crop protection in the UK. Possible applications include most ecological monitoring and surveillance activities, but also the weather or pollution monitoring.

## 1 Introduction

Monitoring networks capture spatiotemporal variations of one or more variables of interest. They offer discrete measurements which can be used to predict the variable at unmeasured times and/or locations (with statistical models, as in *e.g.* Hiemstra et al., 2009), or to investigate the underlying biological or physical processes (with process-based models, as in *e.g.* Roques and Bonnefon, 2016). A desired property of the network is to capture the most significant spatiotemporal variations, so that their subsequent generalisations are meaningful. In this regard, Monitoring Network Optimisation (MNO) is needed to support many surveillance and monitoring activities that are challenged to produce meaningful data from limited resources. Thus, it is imperative that these networks address the optimal selection of sites for the greatest practical and scientific impact.

The question of where and when to sample a process has been approached with design-based and model-based frameworks (Brus and de Gruijter, 1997). The former aims at inferring population parameters while limiting spatial bias, with sampling strategies of varying degrees of randomness and stratification to ensure coverage. The latter requires the development and training of a model of the data generating process to select sampling locations that will reduce the model uncertainty (Mateu and Müller, 2012a), often simply sampling where the model is the most uncertain about its predictions. MNO is a case of optimal sampling which generally builds on an existing network and hence, existing measurements of the variable of interest (Mateu and Müller, 2012b). It is therefore naturally addressed with a model-based approach, in which historic measurements serve the development of a model which subsequently informs site selection.

Optimal sampling designs often rely on modelling approaches such as kriging (Müller, 2007; Spöck, 2012), which builds on the spatial autocorrelation of the measurements and, optionally, their regression to environmental covariates (Oliver and Webster, 2015). However, autocorrelation can appear inconsistent at the scale of sampling, preventing variograms from being fitted, particularly when a small number of measurements are available. This also affects our ability to disentangle complex relationships between measurements and environmental covariates. Machine learning approaches, such as random forests, are usually less hindered by aforementioned conditions (Holloway et al., 2018; Fouedjio and Klump, 2019). A key limitation, however, is that machine learning methods are often unable to associate their predictions with uncertainty estimates. Nonetheless, when uncertainty estimates are provided, by mapping them from the model’s feature space to the geographical space, we can guide the selection of the next sensor location. Using machine learning for such model-based optimal sampling has been referred to as *uncertainty sampling,* an active learning strategy (Tuia et al., 2012). Its principle is iterative: sample where the model uncertainty is the highest, increment the dataset with the new measurement, and train the model again. In our practical case, by deploying sensors in high uncertainty areas, we ensure a strategic increment to the dataset for the next training iteration.

Monitoring network optimisation has been applied to various environmental concerns such as weather (Heuvelink et al., 2012), air pollution (Helle and Pebesma, 2012) or water pollution (Do et al., 2012). However, the question of where and when to sample is also routinely considered by ecologists who are often confronted with domestic and practical concerns that dominate their decision process. These include, but are not limited to, the permission to place monitoring infrastructure in a particular location, the available staff in close proximity to service the infrastructure and the local expertise to identify species to inform measures of occupancy, phenology and abundance collected from the device. These concerns cloud strategic thinking of where a trap is best placed to inform occupancy, phenology and abundance from a biological perspective. Supporting such decision with both statistical evidence and a range of options then allows for an objective, yet flexible improvement.

Our focus is on the Rothamsted Insect Survey (RIS) network of 12.2m suction-traps. These traps are mostly located in agricultural areas and have been continuously monitoring insect migration and movement since 1964 (Bell et al., 2015). For the purpose of this study, the network has two objectives centred around species level prediction of aphid migration events. The first *practical* objective is the efficient and timely detection of seasonal aphid migration, which signals the need for bulletins and alerts to be sent to growers and other interested bodies. With this information, growers have enough time to inspect their crops and respond appropriately, thereby reducing the prophylactic use of insecticides. The second *epistemic* objective is to provide the best representation of aphid migration phenologies in order to increment scientific knowledge through hypotheses testing (Bell et al., 2012, 2015, 2019). This second objective is directly addressed by uncertainty sampling, *i.e.* the network improvement is maximised by setting a trap where the model predictions are the most uncertain. The first *practical* objective however, is about handling a risk: the risk of missing a seasonal infestation, potentially resulting in costly damage on crops. Hence, addressing this objective requires accounting for the distribution of crops locally at risk of infestation.

Since aphids are identified to the species level, we develop a model predicting locally when an aphid species starts its seasonal migration (*i.e.* first flight), thereby signalling when aphids may present a threat to agriculture. This is done for the 14 most common aphid species in the UK, taking the form of a multi-output regression (Borchani et al., 2015). It builds on an artificial neural network (ANN), an architecture enabling the development of complex relationships between inputs (the features, or covariates) and outputs (the labels, or response variables), through a succession of hidden layers of neurons. Because of their unmatched learning abilities, ANNs are proven assets in addressing tasks of ecological complexity (see *e.g.* Ye et al., 2019). Here, for the purpose of uncertainty sampling, we detail how ANNs can associate their predictions with uncertainty estimates. We explain how the *practical* and *epistemic* objectives described above relate to two different sources of uncertainty, both approachable with ANNs through specific augmentations. Finally, building on the estimation of those uncertainties, we provide propositions of improvement for the RIS suction trap network that address both objectives.

## 2 Theory: uncertainty estimation

The promises of uncertainty sampling follow a simple heuristic: by sampling where the model uncertainty is the highest, we maximise the model improvement while limiting the cost of additional sampling (Lewis and Gale, 1994). However, this hides a crucial complexity: a model can be uncertain by lack of consensus in its measurements, or simply by lack of measurement (Kendall and Gal, 2017). The first case defines the *aleatory* uncertainty, the intrinsic variation in the data generating process. It is sometimes called irreducible uncertainty as it is not expected to reduce when generating more samples in a given region of the input space (but could be reduced by expanding the input space with additional features). The second case defines the *epistemic* uncertainty which can be reduced with more samples and represents our lack of knowledge of the data generating process. Note that aleatory and epistemic uncertainties are sometimes referred to as risk and uncertainty respectively (see *e.g.* Osband, 2016).

Which uncertainty to sample depends on the objective of the network. What we previously described as the *epistemic* objective of the network, *i.e.* incrementing our knowledge of the aphid phenology (with *e.g.* better population parameter estimates), should focus on the *epistemic* uncertainty (Nguyen et al., 2019). This means sampling in priority unexplored areas of the input space.

The *practical* objective of the network, *i.e.* the timely detection of aphid arrival, is more complex to address. On one hand, better knowledge of the process means a better generalisation and hence, more reliable declarations of infestation farther away from the traps. On the other hand, by setting traps in areas with greater aleatory uncertainty (*i.e.* where large variations of the response variable are expected), we allow the timely detection in areas where predictions are most unreliable. This second approach provides essentially a local improvement to the network and therefore, the local density of crops to protect should be factored in the decision.

We detail in this section how to estimate those two uncertainties with ANNs (see Hüllermeier and Waegeman, 2020, for a wider perspective on their estimation in machine learning).

### 2.1 Aleatory uncertainty

From similar inputs, a model can output a distribution of responses that informs us on the inherent variability within the measured system, this is known as the aleatory uncertainty. A covariate (or feature) can affect the distribution of the response variable (or label) without affecting its mean value. Such relationships, although important for risk prediction, are likely missed by a regression to the mean. Quantile regressions identify correlations between covariates and specific quantiles of the response variable distribution (Koenker and Bassett, 1978), allowing a more complete view of their relationship. They offer a perspective on extreme events and are therefore particularly relevant to risk management (Herrera et al., 2018). For example, if modelling a pest arrival, predicting a lower quantile of the response variable tells us how early an infestation can be expected locally with a set degree of confidence. The appeal and singularity of this method are that quantiles, and hence prediction intervals, can be derived without concern of error distribution or variance heterogeneity (Cade and Noon, 2003).

By switching from a regression to the mean to a quantile regression, the ANN is trained to output quantiles of the response variable distribution, typically quantiles 5%, 50% and 95%, instead of the mean. While the input shape remains unchanged, the output becomes a 1-dimensional array of as many quantiles as desired (a simultaneous quantile regression, Tagasovska and Lopez-Paz, 2019). This is done using the so-called *pinball* loss function instead of the quadratic loss function (or L2 loss) used in regressions to the mean. The *pinball* loss, more formally referred to as the asymmetric absolute loss function (Furno and Vistocco, 2018), is given by

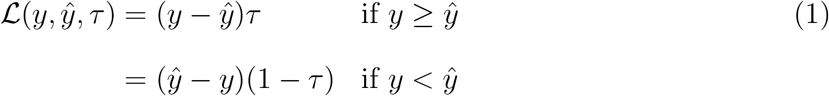

where *y* is an observation, 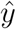 the corresponding prediction and *τ* the desired quantile to estimate.

Quantile regressions are extremely useful for problems with heterogeneous variance across the feature space, such as in our example (and in ecology in general, Cade and Noon, 2003). They can be incorporated in statistical methods such as GAMs (Fasiolo et al., 2017) but are more often seen in machine learning approaches such as random forests (see Quantile RF, Meinshausen, 2006). In contexts of weak spatial autocorrelation of the labels and/or non-linear relationship to the features, quantile random forests have shown more reliable uncertainty estimation than kriging methods (Fouedjio and Klump, 2019). Note also that simultaneous quantile regressions are conveniently supported by the dimensional flexibility of ANNs, whose outputs can easily be made multidimensional and/or multiple (as in our example), and whose generalisation power is improved by the additional information gathered by the multiple quantiles estimated at once (Rodrigues and Pereira, 2018).

### 2.2 Epistemic uncertainty

Epistemic uncertainty marks the lack of exploration of the feature space: the more similar a location is to the sensor locations, the lower its prediction uncertainty. A diversity of methods exist for epistemic uncertainty estimation in ANNs (Gal and Ghahramani, 2016; Gal et al., 2017; Tagasovska and Lopez-Paz, 2019) but *ensemble* methods seem the most accepted (Osband, 2016). Ensemble methods build on a set of models which are trained on various perturbed versions of the data and whose predictions are subsequently averaged (Lakshminarayanan et al., 2017). *Bagging* (from *bootstrap aggregation*, Breiman, 1996) is an ensemble method in which the perturbation results from sampling the training dataset with replacement, *i.e.* bootstrapping. In this approach, the trained ANNs that compose the ensemble are trained on bootstrapped data sets with random weight initialisations, and therefore they output varying predictions. Essentially, bootstrapping allows the approximation of the statistical population by simply resampling the sample data. Therefore, where the predictions of the ensemble disagree the most, the most sensitive to resampling is the approximation, marking the scarcity of data points in the local area of the feature space.

In practice, the output of the bagging ensemble of trained models is an ensemble of predictions, from which both central tendency and dispersion measures are derived. The former constitute more robust predictions than the single model predictions, while the later gives estimates of the epistemic uncertainty. Note that for a quantile regression as in our case, we derive central tendency and dispersion measures of the quantiles of interest.

## 3 Material and Methods

### 3.1 Data

#### 3.1.1 Labels: aphid species arrival

The RIS network initially started with 2 suction traps in 1964 during an experimental period and then formally began reporting with these two traps in 1965. The network grew progressively to 23 traps in 1980 before being reduced to its current size of 15 traps in 1990 (see Figure 1-**A**). Improving the monitoring network requires us to consider the diversity of its measurements and to select those that are key to its optimisation. Therefore, we consider the aphid arrival of the 14 most common species of aphids, trapped between 1965 and 2018 across the UK. These species represent two thirds of the species reported by the RIS on its weekly bulletin (https://insectsurvey.com/).

**Figure 1:**
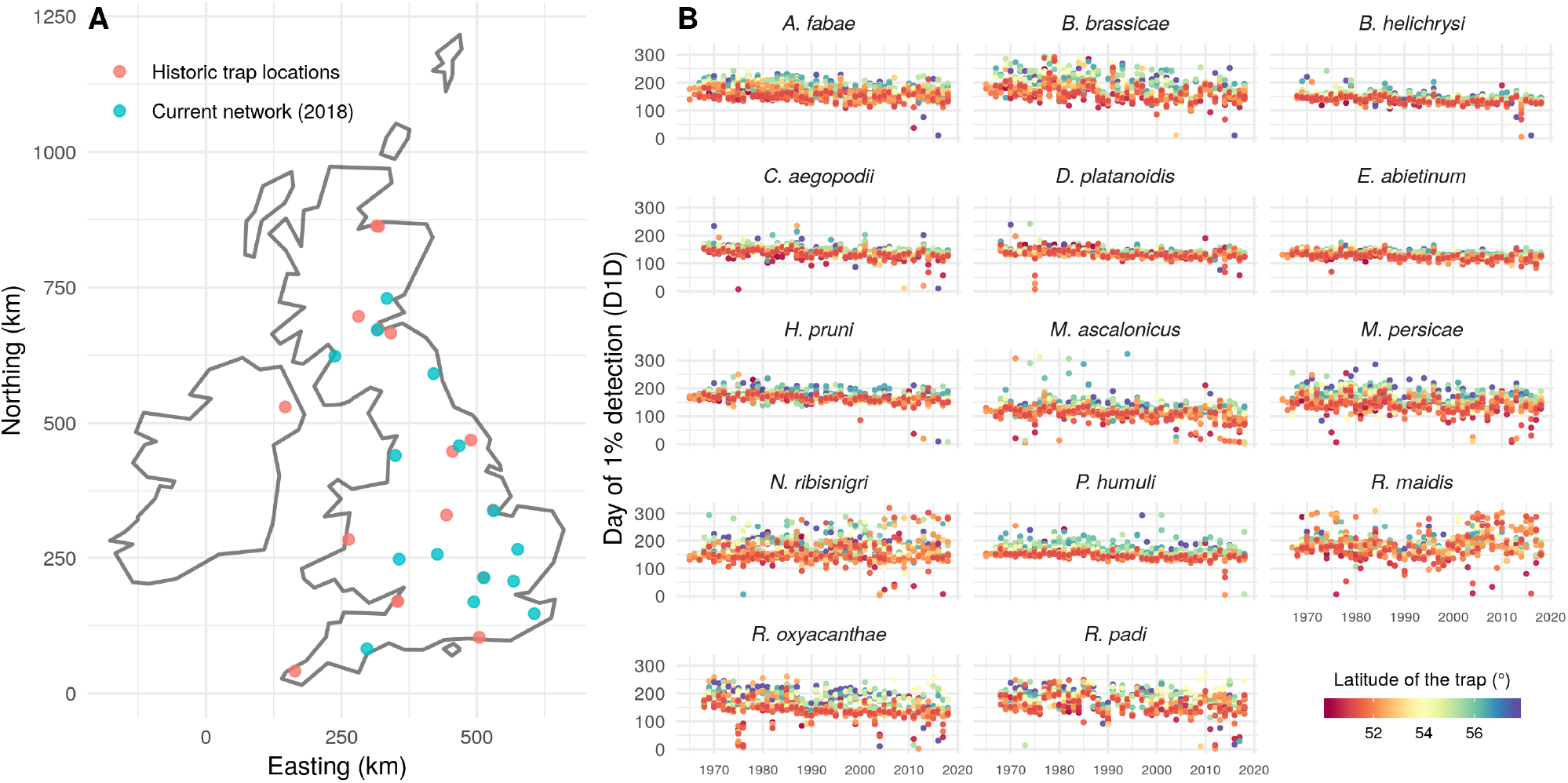
**A** - Historic and current (2018) trap locations across the UK, projected on the British National Grid. **B** - Day of 1% detection of the 14 aphid species measured by the network since 1965.

Aphid arrival happens seasonally and its time of occurrence can be quantified by the day of first flight (DFF). The DFF is a critical phenological event of prime importance for pest control and ecology. It represents the Julian day at which a given species is first detected in a suction-trap catch. However, this variable is subject to large variations caused by the occasional extreme outlier, so predicting instead the day at which a percentile of the yearly catches is reached can be more insightful (as in *e.g.* Thackeray et al., 2010, 2016; Sheppard et al., 2016; Pak et al., 2019). We focused on the day of 1% detection (D1D) (*i.e.* the Julian day that the first percentile of the year abundance was detected), as a proxy for aphid arrival as it is practically equivalent to the DFF and shows a lower variance and a more symmetric distribution, making it easier to predict (see the measured D1D in Figure 1-**B**).

#### 3.1.2 Features: environmental covariates

Aphid phenology, and more precisely aphid arrival, has been shown to depend on climate and land-cover variables. Temperature appears to be the main driver (Zhou et al., 1995) and the current trend for warmer winters translates very distinctly into earlier captures (Harrington et al., 2007; Bell et al., 2019). Figure 1-**B** also illustrates this trend in time, as well as across latitude. Precipitation effects are less direct but also expected through the drought-stress of plant hosts (Mcvean and Dixon, 2001). Finally, land-use also appears a useful predictor, although through non-linear and interacting effects (Cocu et al., 2005b). Therefore, as established drivers of aphid arrival, temperature, precipitation and land-use constitute the key features of our regression.

Climate observations were obtained from the HadUK grid (Hollis et al., 2019). It offers, among many other measurements, daily temperatures and rainfall at varying spatial resolutions (we use 5km × 5km). We simplify the dataset by averaging the daily temperature and rainfall into monthly features, from September of year *n* – 1 to August of year *n*. By doing so we aim to reduce the input noise, as we expect the impact of the weather on aphid phenology to develop over longer time-scales than daily. We also include the number of frost days per winter months.

Land-use data comes from the [CORINE](https://land.copernicus.eu/pan-european/corine-land-cover) land-cover (CLC 2000, 2006, 2012 and 2018) dataset from Copernicus, the EU Earth Observation Programme. This very high spatial resolution dataset (100m×100m) is aggregated into a compositional dataset at the 5km×5km working resolution. Only the coarse and intermediate levels of classification where used, each splitting land-covers into 5 and 15 classes respectively. For both levels, a local measure of the Shannon diversity index was derived and added as covariate to capture the landscape patchwork.

We also added static topographical features to the set of covariates. First, distance to traps, which allows for proximity effects to arise in a more flexible way than by fitting a variogram (Hengl et al., 2018), allowing predictions to build on the spatial autocorrelation of the labels (Cocu et al., 2005a). Finally, elevation and distance to sea were included to capture additional details of the local climates (specifically in regard of wind). The result is a total of 86 features collected everywhere in the UK on a 5km×5km resolution, every year since 1965. Their distributions are figured in Appendix B.

### 3.2 Multi-output quantile regression

Our multi-output quantile regression predicts quantiles of the D1D of the 14 species of interest. It builds on a specific ANN architecture: 3 hidden layers common to the 14 species, followed by one species-specific hidden layer of a smaller size. Complex interactions of covariates can develop in the common layers, while species-specific reactions to those develop in the species-specific layers. This architecture allows flexibility while limiting model complexity (*i.e.* the number of parameters) which greatly improves generalisation by reducing overfitting (Reyes and Ventura, 2019).

Uncertainty estimation is not a common feature in ANN libraries. As mentionned previously, estimating aleatory uncertainty requires the implementation of the specific *pinball* loss function, while, as presented here, estimating epistemic uncertainty requires training an ensemble of networks on bootstrapped data sets (*i.e.* bagging), which is computationally demanding. In addition to those technical developments, and following Osband et al. (2018), we add *Randomised Prior Functions* to the network. This mechanism builds on an untrainable parallel network whose output is added to the trained network output, thereby acting as a Bayesian prior (see Figure 6 in Appendix A for a schematic view of the network). Then, the bagging predictions provide an approximate posterior distribution from which to estimate epistemic uncertainty (Osband et al., 2016). This (technically optional) prior mechanism corrects a tendency for overconfidence in the bootstrap ensemble models in situation of weak measurement heterogeneity (*i.e.* low aleatory uncertainty). Hence, they help disentangling the confusing effects of data scarcity and data heterogeneity, therefore resulting into epistemic uncertainty estimates that remain unaffected by measurements heterogeneity (from which arises aleatory uncertainty, see appendix C.5 of Osband et al., 2018, for an illustration).

In order to reduce the input noise we run a simple feature selection consisting of dropping correlated features with a threshold of 95% correlation. Of the 86 features described above, 63 remained for training. Thresholds of 90% or 100% (no dropping) were also tested but were found to reduce performances. This approach does not guarantee that the removed features are not determinant predictors, it only ensures that the signal they carry is not withdrawn with them as the data set is reduced.

The ANN was developed with Keras, Tensorflow and Scikit-learn in Python 3.6. Details on the ANN architecture, activation, regularisation, model selection, training and residuals are given in Appendix A.

### 3.3 Monitoring network optimisation

To the best of our knowledge, uncertainty sampling is always addressed from an epistemic point of view, aiming at the exploration of the feature space. If, like ordinary kriging, a model builds on the labels’ spatial autocorrelation only, the feature space and the geographical space are confounded and uncertainty arises with the geographical distance from measurement locations (see *e.g.* Chen et al., 2016). In such cases, uncertainty sampling results in space-filling designs. However, as environmental covariates are added to the model, unexplored areas of the feature space, which hold the highest uncertainty, do not necessarily correspond to unexplored areas of the geographical space (Brus and Heuvelink, 2007). Here however, as most of our environmental covariates exhibit strong spatial autocorrelation, a space-filling design could be expected despite their large number. Addressing our epistemic objective of improving the predictions of aphid arrival, we seek to identify the locations where the function

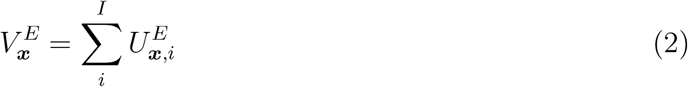

is greatest; where ***x*** marks the spatial coordinates (*x, y*), and 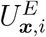 is the epistemic uncertainty of species *i* at location ***x***. As the DFF itself, it is measured in *days.* The measure of *U_E_* used here is the standard error of the lower quantile 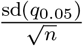, emphasising our focus on early infestation events.

On the other hand, areas of high aleatory uncertainty do not correspond to unexplored regions of the feature space (Nguyen et al., 2019). The predictions there are relatively informed by contemporary and past measurements of the network, but still they remain largely undetermined. Those are areas in which given weather and land-use conditions can produce a wide range of responses, *i.e.* early D1D are likely but so are late ones. In those areas, the nearest trap measurements are poorly informative of aphid arrival, and therefore farmers cannot rely on them for timely reaction. Setting a trap locally would address this issue, provided that such high aleatory uncertainty areas are properly identified by the quantile regression.

However, minimal learning improvement should be expected from setting traps in high aleatory uncertainty areas. Sampling this irreducible uncertainty only addresses the immediate and local objective of timely declaration of infestation. Of course, unpredictable infestations only constitute a risk if susceptible crops are locally present. Hence, to account for the local *value* of a new trap, we started with the distribution of the main crops in the UK (from UKCEH, 2018, [Land Cover® plus](https://www.ceh.ac.uk/crops2015): [Crops maps](https://www.ceh.ac.uk/crops2015), see Figure 2). We then asked aphid experts to devise a link matrix *l_i,j_* between aphid species and the risk they pose to those crops (see Table 1 in Appendix C). Setting a new trap addressing the practical objective can be done by identifying locations where the function

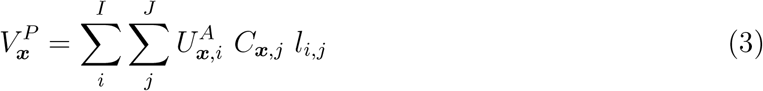

is greatest; here 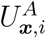 is the aleatory uncertainty of species *i* at location ***x***; *C_x,j_* the proportion of land unit ***x*** occupied by crop *j*; and *l_i,j_* is the susceptibility of crop *j* to species *i*. The measure of *U_A_* used here is the 90% prediction interval of the DFF (with same unit): 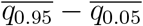.

**Figure 2:**
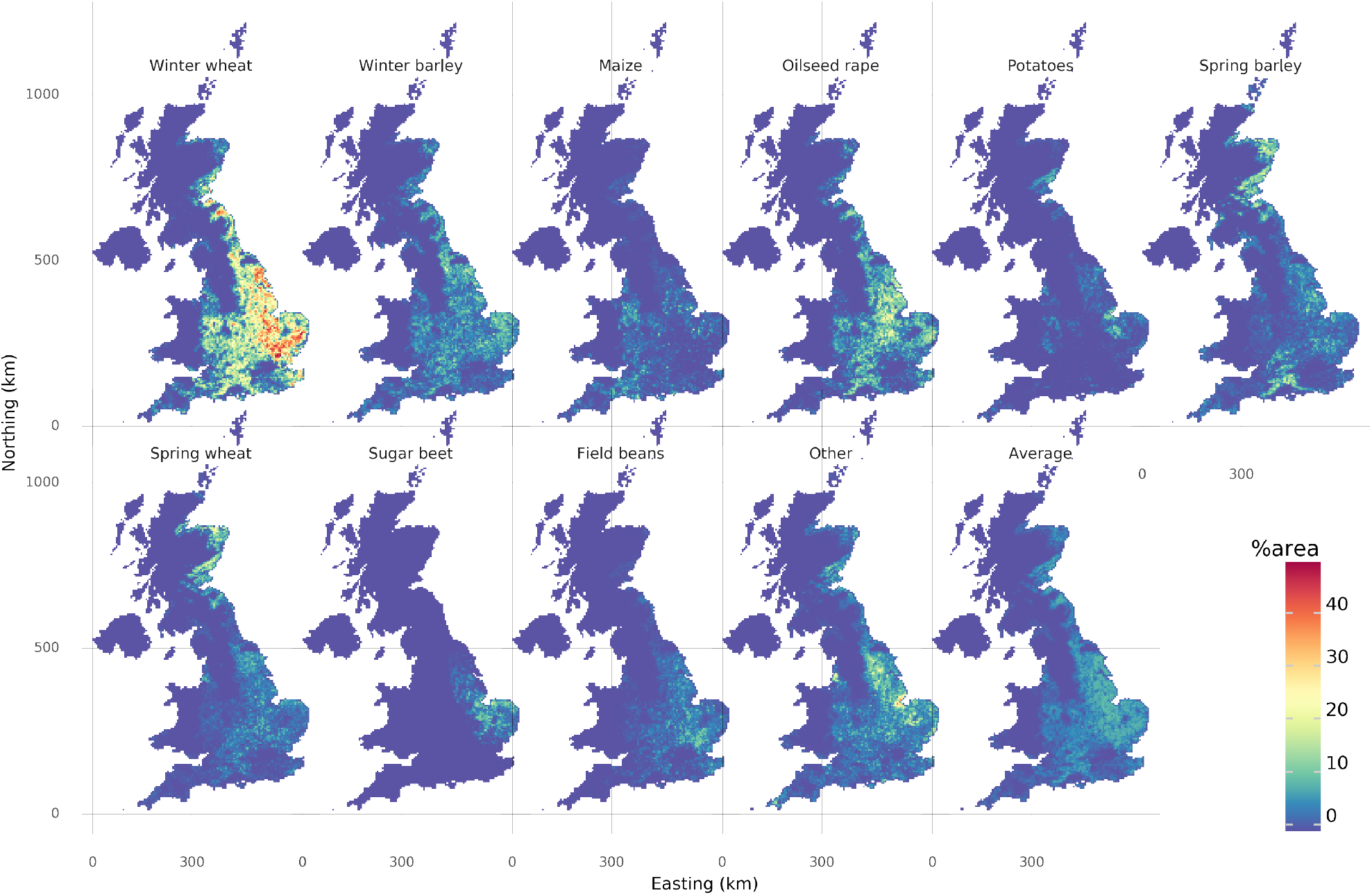
Distribution of the main crops across the UK (missing data for Northern Ireland), projected on the British National Grid.

**Table 1:** Link matrix charactersing crops suceptibilty to the 14 aphid species of interest. Crops here spring barley, spring wheat, winter barley, winter wheat, sugarbeet, fava beans, maize, oilseed rape, potatoes and other (diverse vegtables). A value of 1 marks an established trophic link, a value of 0.5 marks an emerging trophic link, and no values marks the absence of a known trophic link.

| Species | sb | sw | wb | ww | be | fb | ma | or | po | ot | $\mu_{D1D}$ | $\sigma_{D1D}$ |
| --- | --- | --- | --- | --- | --- | --- | --- | --- | --- | --- | --- | --- |
| *M. ascalonicus* | — | — | — | — | — | — | — | 0.5 | 1 | — | 120.0 | 31.6 |
| *R. padi* | 1 | 1 | — | — | — | — | 1 | — | — | — | 173.8 | 35.2 |
| *E. abietinum* | — | — | — | — | — | — | — | — | — | — | 130.9 | 12.6 |
| *D. platanoidis* | — | — | — | — | — | — | — | — | — | — | 136.0 | 16.0 |
| *B. helichrysi* | — | — | 0.5 | — | 0.5 | 1 | — | — | 0.5 | — | 142.3 | 16.6 |
| *C. aegopodii* | — | — | — | — | — | — | — | — | — | — | 141.2 | 19.1 |
| *N. ribisnigri* | — | — | — | — | — | — | — | — | — | — | 164.9 | 40.4 |
| *M. persicae* | 0.5 | 0.5 | 1 | 1 | 1 | 1 | 1 | 1 | 1 | — | 155.9 | 34.4 |
| *R. maidis* | — | — | — | — | — | — | — | — | — | — | 183.6 | 41.0 |
| *H. pruni* | — | — | — | 0.5 | — | — | — | — | — | — | 168.5 | 19.2 |
| *P. humuli* | — | — | — | — | — | — | — | — | — | — | 155.5 | 22.2 |
| *R. oxyacanthae* | — | 0.5 | — | — | — | — | — | — | — | — | 158.2 | 37.4 |
| *A. fabae* | — | — | 1 | 1 | 1 | 1 | 1 | — | 1 | — | 167.8 | 27.2 |
| *B. brassicae* | — | — | — | — | 0.5 | — | — | 1 | — | — | 176.0 | 38.9 |

## 4 Results

### 4.1 Epistemic objective: predicting migration

For most aphid species, the epistemic uncertainty of the D1D increases with altitude (see Figure 3). Those areas, in addition to being historically deprived of measurements, are environmentally distinct from most trap locations, mainly because they are colder which likely delays first flights until later in the season. They represent a sharply defined region of the feature space in which predictions are merely extrapolations of distant measurements. It is therefore unsurprising to see that the model predictions are the most uncertain at altitude.

**Figure 3:**
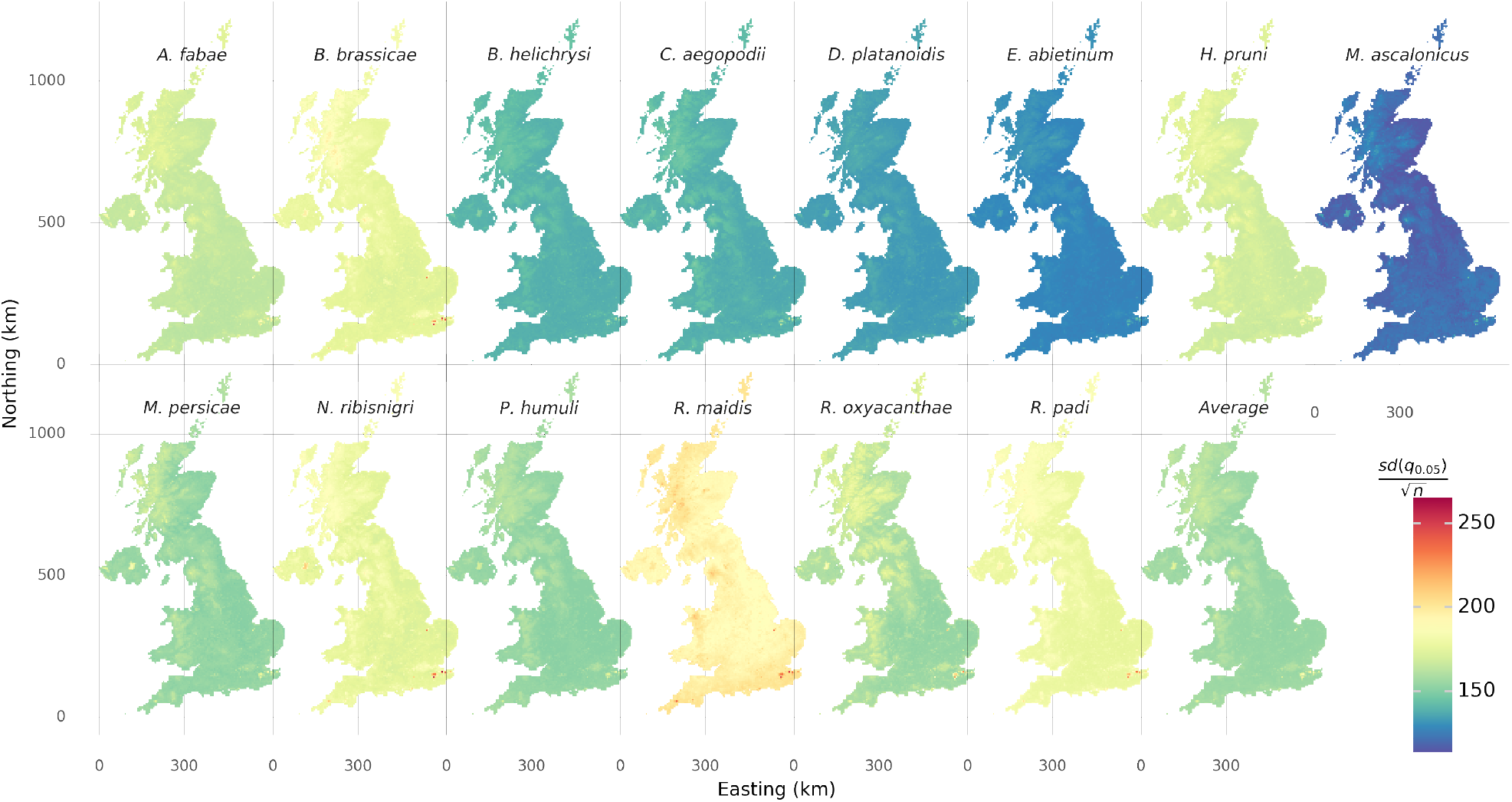
Epistemic uncertainties of the Day of 1% detection for the 14 aphid species. It is expressed as the standard error of the lower quantile: 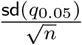 and projected on the British National Grid.

**Figure 4:**
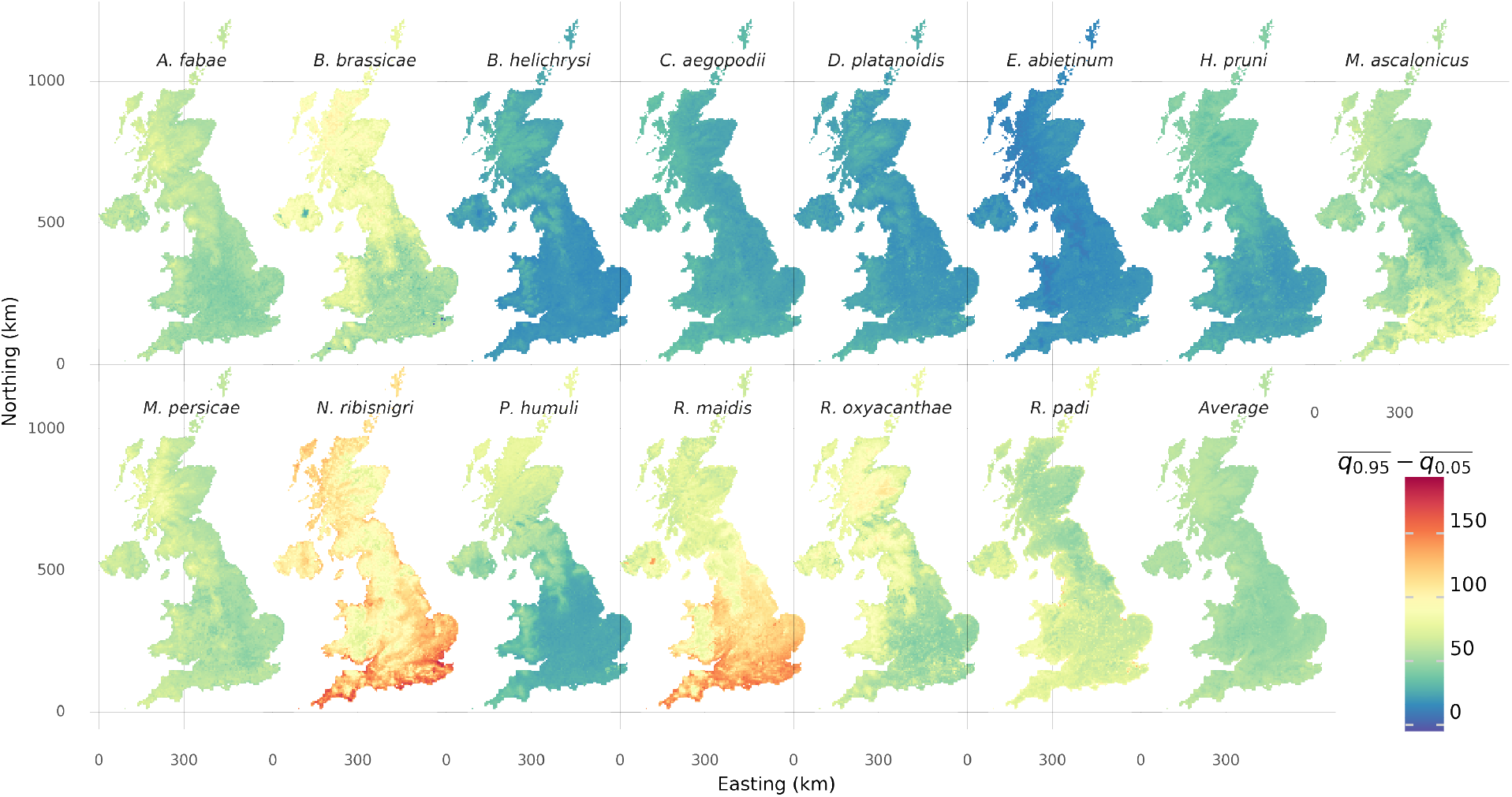
Aleatory uncertainty of the Day of 1% detection for the 14 aphid species. It is expressed as the 90% prediction interval: 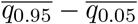 and projected on the British National Grid.

However, setting a trap in the Highlands might seem irrelevant to agricultural production, as in addition to there being no crops to protect there, aphids are not a problem in such environments. It is nonetheless where the best generalisation improvement would occur, maximising the reduction of the overall uncertainty at the next training iteration (*i.e.* after next season catches). However, once we exclude from the problem regions of limited agronomic interest, new areas of high uncertainty appear. Figure 5-**A** shows that Cornwall, Northern Ireland or Dyfed county in Wales are places of high agricultural interest that are not only geographically remote from the current monitoring network, but also environmentally different from it. In other words, new trap locations are not only selected by the spatial imbalance of the current trap locations, but by the sampling imbalance within the whole feature space.

**Figure 5:**
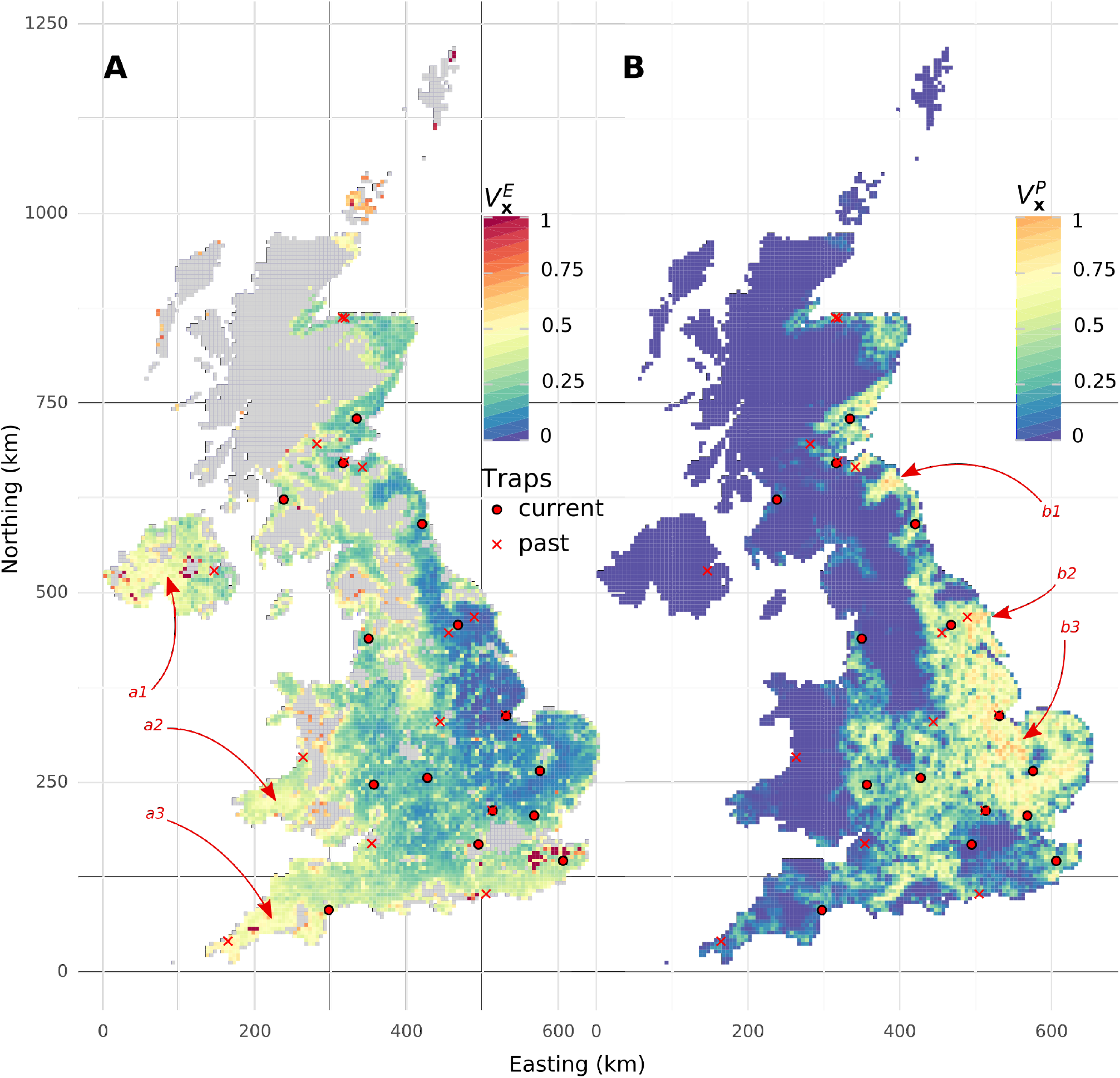
**A** - Standardised value of a new trap in addressing the epistemic objective following Equation 2 (in areas where at least 1/3 of the land serves an agricultural purpose, including pasture). **B** - Standardised value of a new trap in addressing the practical objective following Equation 3 (missing Northern Ireland data). The red arrows show propositions of new trap locations answering best the epistemic objective: in Northern Ireland (a1), in the Dyfed county (a2) or in Cornwall (a3); and the practical objective: near the river Tweed estuary (b1), in Yorkshire (b2) or in Cambridgeshire (b3). Projection on the British National Grid.

### 4.2 Practical objective: protecting crops

Figure 4 shows the distribution of aleatory uncertainty of the D1D for each species. Note that it is distributed differently to the epistemic uncertainty (Figure 3). From the information in Figures 4 and 2, and according to Equation 3, optimal locations for a new trap can be derived (see Figure 5-**B**). Several high value areas are highlighted, notably along the river Tweed on the English-Scottish border, in Cambridgeshire, or also in Yorkshire where as shown in Figure 5-**B** a trap used to be operational.

## 5 Discussion

Through a logical process of estimating aphid phenology relative to meteorological and landscape drivers, our models show that critical areas of agricultural importance are lacking the trap infrastructure needed to estimate first flight and other metrics. If populated, these new trap locations will improve decision support in UK agriculture, thereby reducing risk of deleterious feeding by aphids and virus transmission by aphids to crops.

The network can be improved in two ways, for improved prediction of aphid arrival (the epistemic objective), or for reduced risk of missing crop infestations (the practical objective). The former is a UK-wide improvement, while the latter is a local improvement that depends on the presence of crops needing protection. For both, our approach proposes multiple locations for setting a new trap in, providing the decision maker with the degrees of freedom needed in the process of augmenting the current infrastructure. To address this, we have relied on ANNs flexibility.

Although they are mostly known for classification problems, ANNs have long proven their ability for non-linear and multivariate regressions (see *e.g.* Comrie, 1997). For the specific MNO problem addressed here, the basic feed-forward ANN needed to be augmented in 3 ways: (1) become a *multi-output regression* to account for the 14 species of interest, (2) become a *quantile regression* so that aleatory uncertainty could be examined, as well as (3) be fitted *randomised prior functions* and trained in a *bagging* approach for epistemic uncertainty estimation. The results comprise UK-wide predictions of the date of aphid arrival (specifically the D1D) of the 14 aphid species of interest, associated with estimates of the aleatory and epistemic uncertainties.

Uncertainty sampling promises a maximised model improvement by pointing out where additional measurements are the most needed, *i.e.* where the model’s predictions are the most uncertain. Although simple in principle, it requires the distinction of uncertainty sources. It’s target is typically exclusively the epistemic uncertainty (Nguyen et al., 2019). However, many monitoring networks, like the present one, are also destined to detect and prevent a risk, which is better approached by aleatory uncertainty. We showed that those two uncertainties distribute differently, resulting in diverging propositions of location for setting a new trap. Proceeding further is then a subjective choice, balancing risk management and a quest for knowledge.

Uncertainty sampling normally proceeds iteratively, presenting an *oracle* (*e.g.* a human operator) with an uncertain sample to label, which is subsequently added to the training dataset, enabling deeper learning at the next training iteration. The MNO problem presented here is a single iteration of this active learning process, in which the *oracle*’s answer results from setting a new trap for the next season measurement. From season to season, as a slow-paced active learning process unfolds, the learner (*i.e.* our regression model) improves.

The network improvement discussed here is based on the simplest lever: adding a new trap. However, we could address the MNO from a more comprehensive perspective, including addition, deletion and relocation of one or more traps. Finding the optimal scenario would then require accounting for costs, both of setting a new trap and of loosing a given crop to aphids. Because of the scales at stake, such an experiment should be done *in silico*, *e.g.* training a process-based model of the aphid population dynamics, and optimising an iterative improvement process. This would help in appreciating the potential improvement offered by a single trap addition.

In practice, many factors other than reducing uncertainty or risk affect decisions about network improvement. For example, we proposed here that the epistemic objective should account for the presence of agricultural areas, blurring already the distinction with the practical risk-based objective. It is also important to mention that sensors have perception ranges within which a new sensor is merely redundant (a radius estimated to 80km for the suction traps, Benton et al., 2002). Accounting for such an exclusion radius will mechanically result in more space-filling designs. Additionally, the domestic and practical considerations mentioned in the Introduction, such as having staff able to regularly visit traps and identify catches, are also constitutive of such a decision process. Nonetheless, our approach is not scale, location or infrastructure dependent and could be used to improve most ecological monitoring and surveillance activities across the world that have specific network objectives, such as detection of occupancy, phenology and abundance.

As training ANN requires large data sets, our 881 samples appear a limitation of this study. However, every sample includes 14 response measurements and, as they are used to train mostly shared parameters, the small data set effect is attenuated. Nonetheless, it is clear that simpler machine learning methods would provide equivalent learning to this exemplary case. However, only ANNs provide all the flexibility needed here, with custom loss functions, multi-outputs and bagging. This makes ANNs great assets for uncertainty estimation even with small data sets, provided that overfitting is properly controlled.

An immediate extension to this work is therefore its application to rest of Europe (as did Harrington et al., 2007, also with suction traps). The incremented dataset would satisfy the inherent *data-hungriness* of ANNs, while also extending the input space to new climates. In the UK, this would result in improved predictions, and in different epistemic uncertainty distributions as a lot of predictions would no longer result from extrapolation.

## Reproducibility

The ANN training and predictions can be reproduced from the publicly available online [dataset](https://repository.rothamsted.ac.uk/item/979yv/aphid-arrival-in-the-uk-from-1965-to-2018-measured-by-the-ris-suction-trap-network) (Bourhis et al., 2020) and [code](https://gitlab.com/Yo-B/monitoring-network-optimisation/-/tree/master).

## Acknowledgements

The work at Rothamsted forms part of the Smart Crop Protection (SCP) strategic programme (BBS/OS/CP/000001) funded through the Biotechnology and Biological Sciences Research Council (BBSRC) Industrial Strategy Challenge Fund. The Rothamsted Insect Survey, a National Capability, is funded by the BBSRC under the Core Capability Grant BBS/E/C/000J0200. The authors are also grateful to the Alex Dye, aphidologist at Rothamsted research, for devising the link matrix between crops and aphid species.

## Appendices

### A Model architecture, selection and residuals

Our artificial neural network architecture comprises two dichotomies. Firstly, there is a shared part of the network and species-specific parts. The former is detailed in the upper half of Figure 6, while the bottom part explains the latter. Secondly, there is a trained part of the network and an untrained one (in grey in Figure 6).

**Figure 6:**
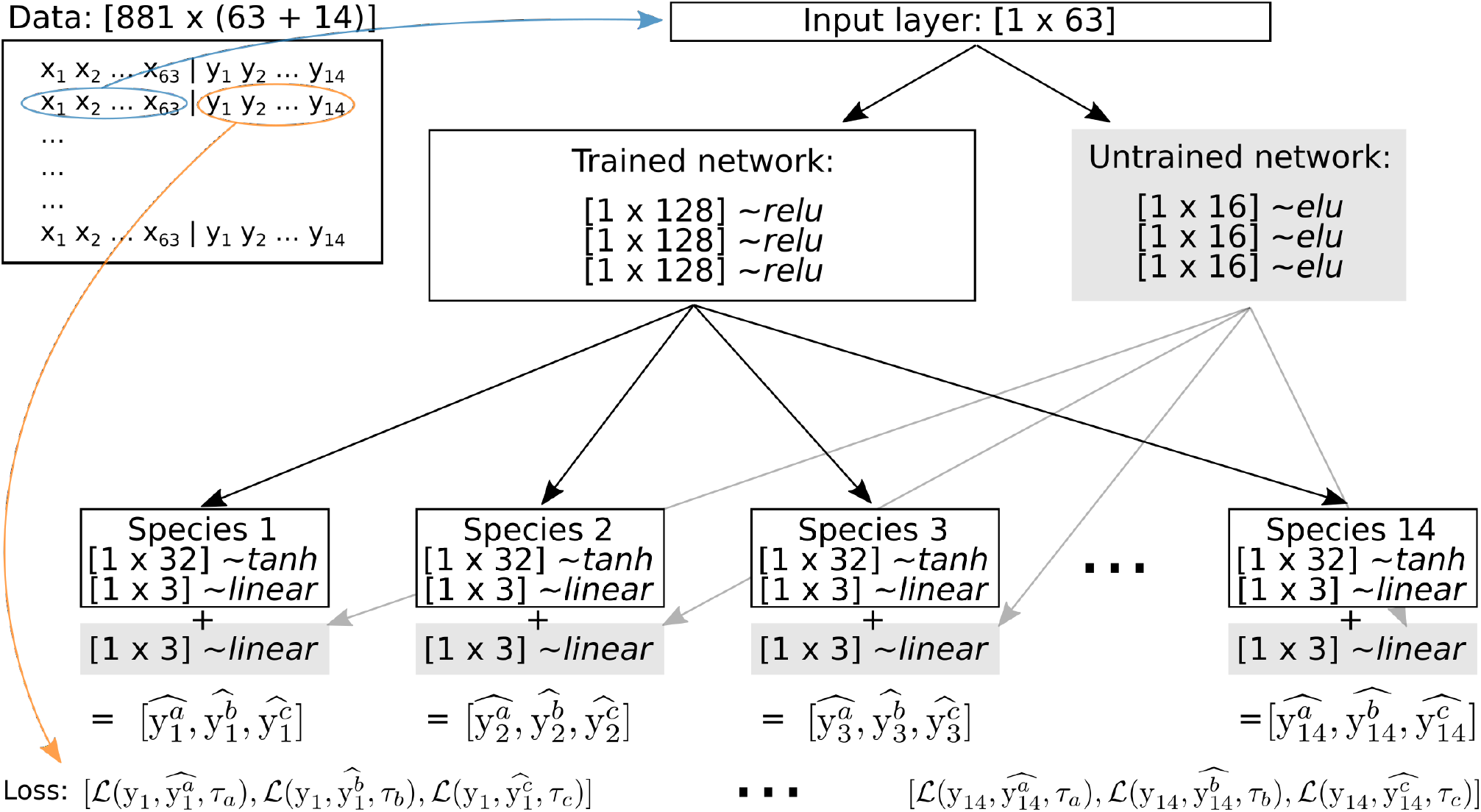
Schematic view of the network architecture, layers dimensions, activation functions, inputs and outputs. 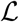 is the pinball loss function given in Equation 1, with *τ_a_, τ_b_* and *τ_c_* mark the target quantiles.

The data is composed of 881 samples for which we have 63 features (the environmental covariates), noted *x*_1_ to *x*_63_, and 14 labels (the D1D of the 14 species of interest), noted *y*_1_ to *y*_14_. The 14 labels could be output as a 1-dimensional array, but as the different quantiles are already output this way, the 14 labels require multi-output. We therefore have 14 outputs, each one of length 3 for the 3 quantiles of interest: 5%, 50% and 95%.

Various activation functions are used here. The output activation function is *linear* as is common for regression. The species specific layers are activated with *tanh* functions, while the main shared network has *ReLU*, rectified linear units, activation. The untrained shared network is activated with *elu*, exponential linear units, functions (Michelucci, 2018).

The network deepness (*i.e.* number of successive layers), the layers dimensions (*i.e.* number of units per layer), as well as as well as, subsequently, the ratio of shared/species-specific number of parameters are hyper-parameters requiring tuning. Tuning, or more formally model selection, was done through cross-validation, reserving 20% of the samples, the testing dataset, and using the 80% remaining for training. The split is done according to a semi-deterministic method referred to as *systemic stratified* (Wu et al., 2013), shown to best distribute the samples when applied to small data sets and low label variance (Zheng et al., 2018), which is our case here. This procedure of random splitting and subsequent training/testing is iterated 10 times, allowing model selection to be based on the averaged testing performance.

Model selection was conducted with regard to the root mean square error (RMSE) between the predicted 50*^th^* percentile of the D1D and the observed D1D. ANNs were instantiated according to a complete factorial design crossing the following hyper-parameters: *number of shared layer* ∈ {1,2, 3,4}, *shared layer size* ∈ {32,64,128,192} and *species-specific layer size* ∈ {0,16, 32,48}. Training was done with the Adam optimiser, with a batch size of 10 samples, and a patience-based callback of 50 epochs without improvement as stopping criterion.

Figure 7 shows no performance improvement beyond 100000 parameters. It also shows that many different designs provide equivalent performances. The retained architecture, as illustrated by Figure 6, is composed of 3 shared layers of 128 units, followed by a set of species-specific layer of 32 units.

**Figure 7:**
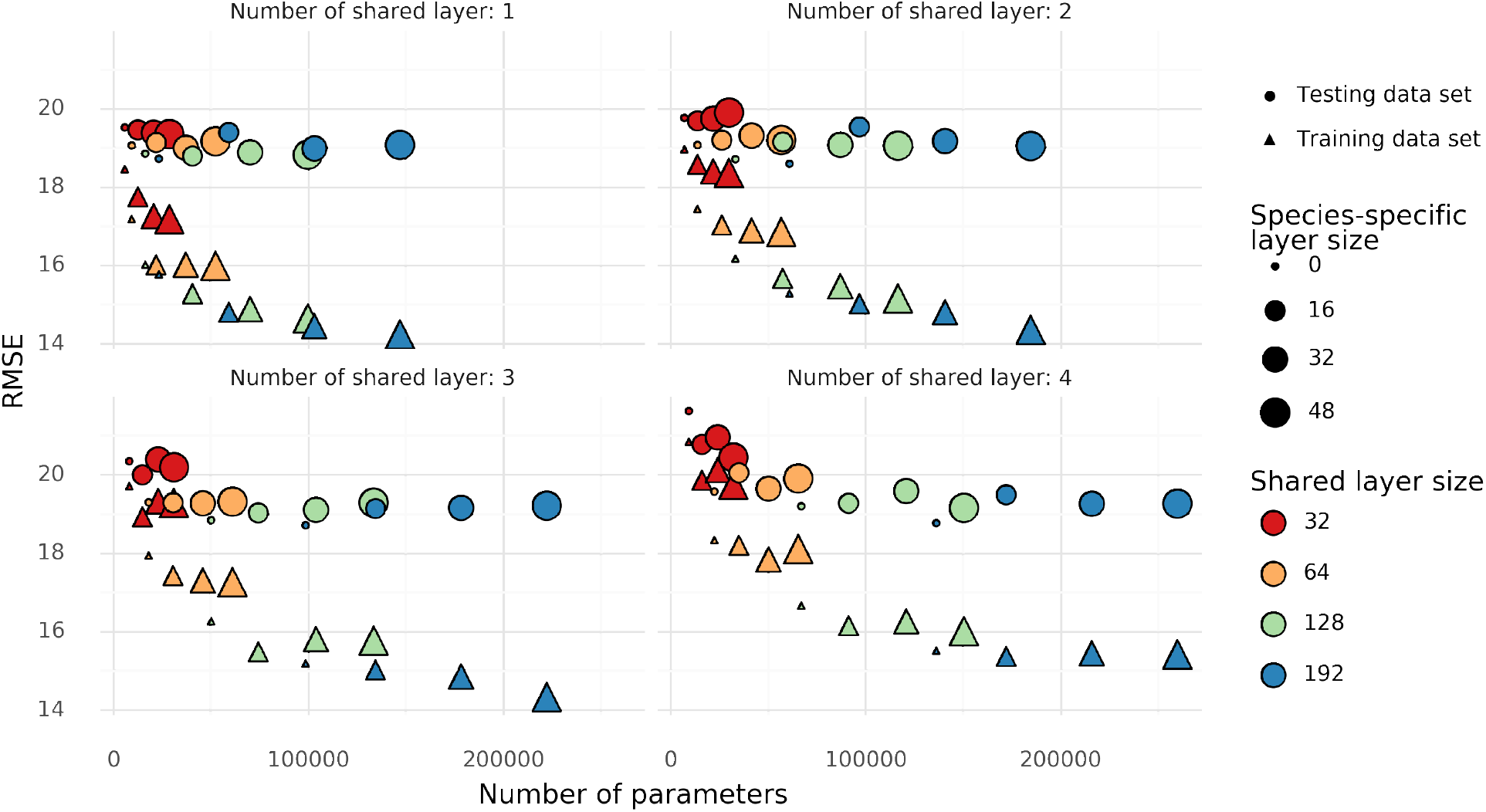
Average RMSE reached by ANNs of varying complexity. Performance on testing data set (the circles) is the key to the model selection procedure.

Figure 8 shows the per-species performances of the retained model. The averaged RMSE results from the cross-validation, in which samples are sequentially part of the training and testing datasets.

**Figure 8:**
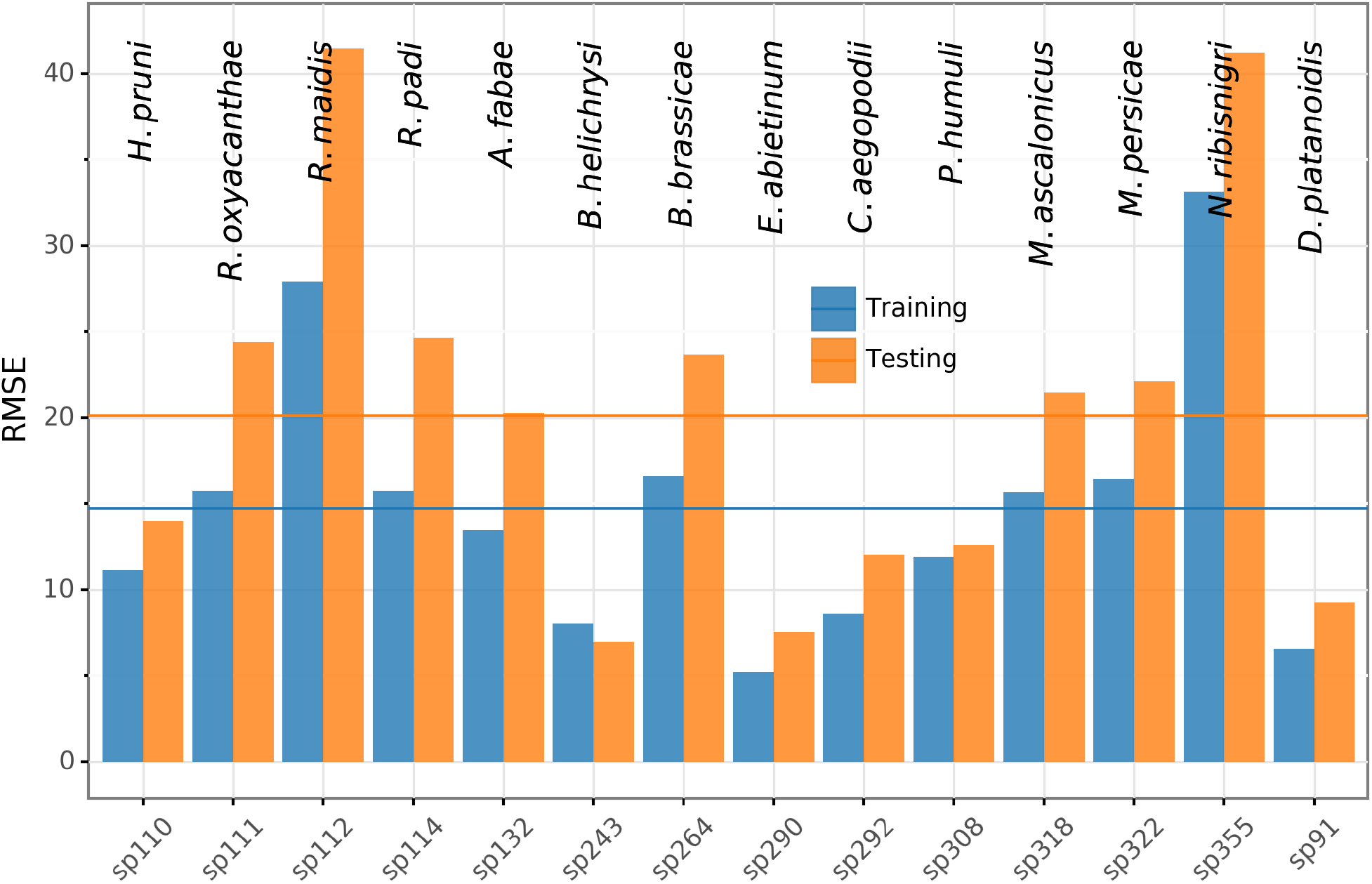
Averaged training and testing RMSE reached by the retained ANN for the 14 different aphid species.

Following model selection, training is done within a bagging procedure. Figure 9 shows the bagging residuals (*i.e.* averaged outputs) for every aphid species as predicted by the selected model. For the sake of illustration, here the dataset is also split into testing and training datasets. However, the whole dataset is used for training once the objective is to map uncertainty across the UK (Figures 3 and 4).

**Figure 9:**
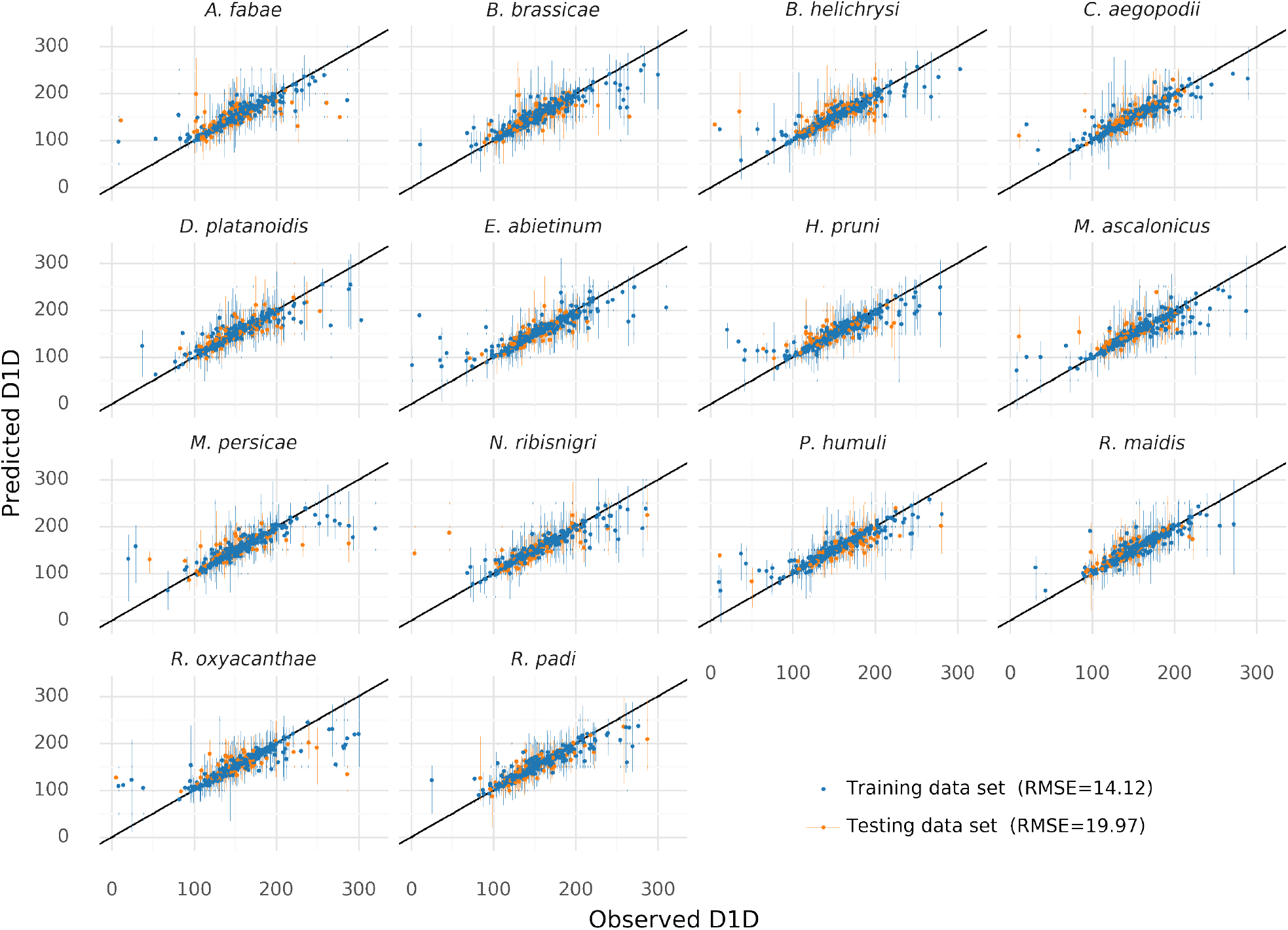
Residuals, *i.e.* predicted vs observed D1D, for the 14 aphid species of interest. This is the output of a bagging ensemble of 30 trained ANNs. For illustration, tThe dataset is split into a training set (in blue, 4/5 of the samples which get bootstrapped in the bagging procedure) and a testing set (in orange, 1/5 of the samples). On the predicted axis, the error bars are built on the average 5% and 95% quantiles, while the predicted point is the average 50% quantile.

**Figure 10:**
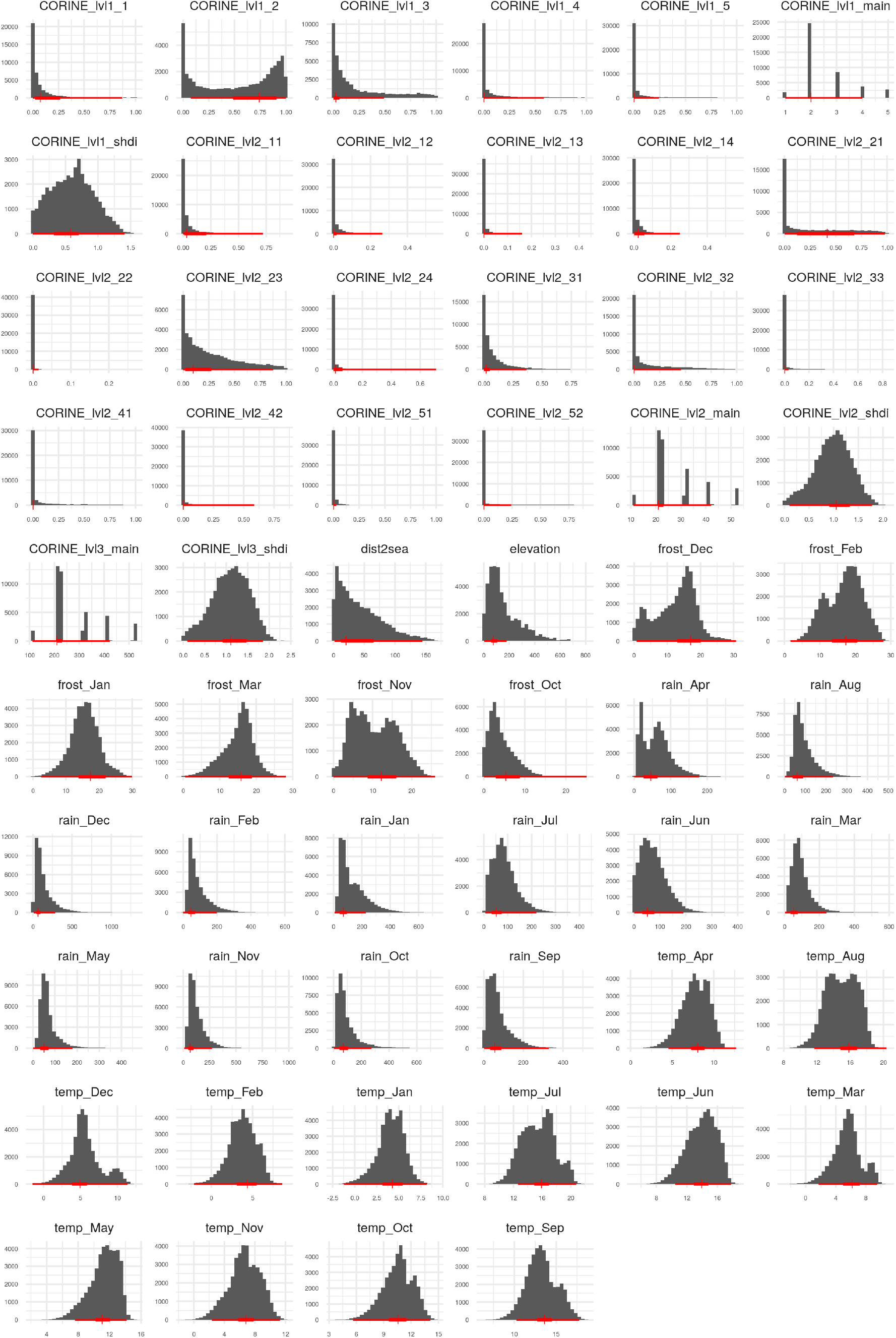
Distributions of the features (except buffer distances) used for regression. The histograms show the distribution of values across the whole UK grid since 1965, while the red lines illustrates the values found at trap locations *(i.e.* used for training) in 5 values: minimum, maximum, 25*^th^*, 50*^th^* and 75*^th^* percentiles.

### B Features distributions

### C Link matrix

## References

Bell, J. R., Alderson, L., Izera, D., Kruger, T., Parker, S., Pickup, J., Shortall, C. R., Taylor, M. S., Verrier, P., and Harrington, R. (2015). Long-term phenological trends, species accumulation rates, aphid traits and climate: five decades of change in migrating aphids. Journal of Animal Ecology, 84(1):21–34.

Bell, J. R., Botham, M. S., Henrys, P. A., Leech, D. I., Pierce-Higgins, J. W., Shortall, C. R., Brereton, T. M., Pickup, J., and Thackeray, S. J. (2019). Spatial and habitat variation in aphid, butterfly, moth and bird phenologies over the last half century. Global Change Biology.

Bell, J. R., Taylor, M. S., Shortall, C. R., Welham, S. J., and Harrington, R. (2012). The trait and host plant ecology of aphids and their distribution and abundance in the United Kingdom: Aphid trait ecology. Global Ecology and Biogeography, 21(4):405–415.

Benton, T. G., Bryant, D. M., Cole, L., and Crick, H. Q. P. (2002). Linking agricultural practice to insect and bird populations: a historical study over three decades. Journal of Applied Ecology, 39(4):673–687.

Borchani, H., Varando, G., Bielza, C., and Larrañaga, P. (2015). A survey on multi-output regression: Multi-output regression survey. Wiley Interdisciplinary Reviews: Data Mining and Knowledge Discovery, 5(5):216–233.

Bourhis, Y., Bell, J., Vandenbosch, F., and Milne, A. (2020). Aphid arrival in the UK from 1965 to 2018 measured by the RIS suction-trap network. Publisher: Rothamsted Research.

Breiman, L. (1996). Bagging Predictors. Machine Learning, 24(2):123–140.

Brus, D. J. and de Gruijter, J. J. (1997). Random sampling or geostatistical modelling? Choosing between design-based and model-based sampling strategies for soil (with discussion). Geoderma, 80(1):1–44.

Brus, D. J. and Heuvelink, G. B. (2007). Optimization of sample patterns for universal kriging of environmental variables. Geoderma, 138(1-2):86–95.

Cade, B. S. and Noon, B. R. (2003). A gentle introduction to quantile regression for ecologists. Frontiers in Ecology and Environment, pages 412–420.

Chen, K., Ni, M., Cai, M., Wang, J., Huang, D., Chen, H., Wang, X., and Liu, M. (2016). Optimization of a Coastal Environmental Monitoring Network Based on the Kriging Method: A Case Study of Quanzhou Bay, China. BioMed Research International. ISSN: 2314-6133 Library Catalog: www.hindawi.com Pages: e7137310 Publisher: Hindawi Volume: 2016.

Cocu, N., Harrington, R., Hullé, M., and Rounsevell, M. D. A. (2005a). Spatial autocorrelation as a tool for identifying the geographical patterns of aphid annual abundance. Agricultural and Forest Entomology, 7(1):31–43.

Cocu, N., Harrington, R., Rounsevell, M. D. A., Worner, S. P., Hullé, M., and the EXAMINE project participants (2005b). Geographical location, climate and land use influences on the phenology and numbers of the aphid, Myzus persicae, in Europe: Environmental influences on aphid distribution. Journal of Biogeography, 32(4):615–632.

Comrie, A. C. (1997). Comparing Neural Networks and Regression Models for Ozone Forecasting. Journal of the Air & Waste Management Association, 47(6):653–663. Publisher: Taylor & Francis _eprint: https://doi.org/10.1080/10473289.1997.10463925.

Do, H. T., Lo, S.-L., Chiueh, P.-T., and Phan Thi, L. A. (2012). Design of sampling locations for mountainous river monitoring. Environmental Modelling & Software, 27-28:62–70.

Fasiolo, M., Goude, Y., Nedellec, R., and Wood, S. N. (2017). Fast calibrated additive quantile regression. arXiv:1707.03307 [stat]. arXiv: 1707.03307.

Fouedjio, F. and Klump, J. (2019). Exploring prediction uncertainty of spatial data in geostatistical and machine learning approaches. Environmental Earth Sciences, 78(1):38.

Furno, M. and Vistocco, D. (2018). Quantile Regression: Estimation and Simulation. John Wiley & Sons. Google-Books-ID: DJhlDwAAQBAJ.

Gal, Y. and Ghahramani, Z. (2016). Dropout as a Bayesian Approximation: Representing Model Uncertainty in Deep Learning. arXiv:1506.02142 [cs, stat]. arXiv: 1506.02142.

Gal, Y., Hron, J., and Kendall, A. (2017). Concrete Dropout. arXiv:1705.07832 [stat]. arXiv: 1705.07832.

Harrington, R., Clark, S. J., Welham, S. J., Verrier, P. J., Denholm, C. H., Hulle, M., Maurice, D., Rounsevell, M. D., Cocu, N., and EUROPEAN UNION EXAMINE CONSORTIUM (2007). Environmental change and the phenology of European aphids. Global Change Biology, 13(8):1550–1564.

Helle, K. B. and Pebesma, E. (2012). Stationary Sampling Designs Based on Plume Simulations. In Spatio-Temporal Design, pages 319–344. John Wiley & Sons, Ltd. Section: 14 _eprint: https://onlinelibrary.wiley.com/doi/pdf/10.1002/9781118441862.ch14.

Hengl, T., Nussbaum, M., Wright, M. N., Heuvelink, G. B. M., and Gräler, B. (2018). Random forest as a generic framework for predictive modeling of spatial and spatio-temporal variables. PeerJ, 6:e5518.

Herrera, M., Ramallo-González, A. P., Eames, M., Ferreira, A. A., and Coley, D. A. (2018). Creating extreme weather time series through a quantile regression ensemble. Environmental Modelling & Software, 110:28–37.

Heuvelink, G. B. M., Griffith, D. A., Hengl, T., and Melles, S. J. (2012). Sampling Design Optimization for Space-Time Kriging. In Spatio-Temporal Design, pages 207–230. John Wiley & Sons, Ltd. Section: 9 _eprint: https://onlinelibrary.wiley.com/doi/pdf/10.1002/9781118441862.ch9.

Hiemstra, P. H., Pebesma, E. J., Twenhöfel, C. J., and Heuvelink, G. B. (2009). Real-time automatic interpolation of ambient gamma dose rates from the dutch radioactivity monitoring network. Computers & Geosciences, 35(8):1711–1721.

Hollis, D., McCarthy, M., Kendon, M., Legg, T., and Simpson, I. (2019). HadUK-Grid—A new UK dataset of gridded climate observations. Geoscience Data Journal, 6(2):151–159.

Holloway, P., Kudenko, D., and Bell, J. R. (2018). Dynamic selection of environmental variables to improve the prediction of aphid phenology: A machine learning approach. Ecological Indicators, 88:512–521.

Huällermeier, E. and Waegeman, W. (2020). Aleatoric and Epistemic Uncertainty in Machine Learning: An Introduction to Concepts and Methods. arXiv:1910.09457 [cs, stat]. arXiv: 1910.09457.

Kendall, A. and Gal, Y. (2017). What uncertainties do we need in bayesian deep learning for computer vision? CoRR, abs/1703.04977.

Koenker, R. and Bassett, G. (1978). Regression Quantiles. Econometrica, 46(1):33.

Lakshminarayanan, B., Pritzel, A., and Blundell, C. (2017). Simple and Scalable Predictive Uncertainty Estimation using Deep Ensembles. arXiv:1612.01474 [cs, stat]. arXiv: 1612.01474.

Lewis, D. D. and Gale, W. A. (1994). A Sequential Algorithm for Training Text Classifiers. arXiv:cmp-lg/9407020. arXiv: cmp-lg/9407020.

Mateu, J. and Müller, W. G. (2012a). Collecting Spatio-Temporal Data. In SpatioTemporal Design, pages 1–36. John Wiley & Sons, Ltd. Section: 1 _eprint: https://onlinelibrary.wiley.com/doi/pdf/10.1002/9781118441862.ch1.

Mateu, J. and Müller, W. G., editors (2012b). Spatio-temporal Design: Advances in Efficient Data Acquisition. Wiley, Chichester, West Sussex, UK, 1 edition edition.

Mcvean, R. I. K. and Dixon, A. F. G. (2001). The effect of plant drought-stress on populations of the pea aphid Acyrthosiphon pisum. Ecological Entomology, 26(4):440–443.

Meinshausen, N. (2006). Quantile Regression Forests. Journal of Machine Learning Research, 7:17.

Michelucci, U. (2018). Applied Deep Learning: A Case-Based Approach to Understanding Deep Neural Networks. Apress. Google-Books-ID: z1ptDwAAQBAJ.

Müller, W. G. (2007). Collecting Spatial Data: Optimum Design of Experiments for Random Fields. Springer Science & Business Media. Google-Books-ID: ivyV9UpL8XkC.

Nguyen, V.-L., Destercke, S., and Huällermeier, E. (2019). Epistemic Uncertainty Sampling. arXiv:1909.00218 [cs, stat]. arXiv: 1909.00218.

Oliver, M. A. and Webster, R. (2015). Basic Steps in Geostatistics: The Variogram and Kriging. SpringerBriefs in Agriculture. Springer International Publishing.

Osband, I. (2016). Risk versus Uncertainty in Deep Learning: Bayes, Bootstrap and the Dangers of Dropout. Workshop on Bayesian Deep Learning, NIPS 2016, Barcelona, Spain., page 5.

Osband, I., Aslanides, J., and Cassirer, A. (2018). Randomized Prior Functions for Deep Reinforcement Learning. arXiv:1806.03335 [cs, stat]. arXiv: 1806.03335.

Osband, I., Blundell, C., Pritzel, A., and Van Roy, B. (2016). Deep Exploration via Bootstrapped DQN. arXiv:1602.04621 [cs, stat]. arXiv: 1602.04621.

Pak, D., Biddinger, D., and Bjørnstad, O. N. (2019). Local and regional climate variables driving spring phenology of tortricid pests: a 36 year study. Ecological Entomology, 44(3):367–379.

Reyes, O. and Ventura, S. (2019). Performing Multi-Target Regression via a Parameter Sharing-Based Deep Network. International Journal of Neural Systems, 29(09):1950014.

Rodrigues, F. and Pereira, F. C. (2018). Beyond expectation: Deep joint mean and quantile regression for spatio-temporal problems. arXiv:1808.08798 [cs, stat]. arXiv: 1808.08798.

Roques, L. and Bonnefon, O. (2016). Modelling population dynamics in realistic landscapes with linear elements: A mechanistic-statistical reaction-diffusion approach. PLOS ONE, 11(3):1–20.

Sheppard, L. W., Bell, J. R., Harrington, R., and Reuman, D. C. (2016). Changes in large-scale climate alter spatial synchrony of aphid pests. Nature Climate Change, 6(6):610–613. Number: 6 Publisher: Nature Publishing Group.

Spöck, G. (2012). Spatial sampling design based on spectral approximations to the random field. Environmental Modelling & Software, 33:48–60.

Tagasovska, N. and Lopez-Paz, D. (2019). Single-Model Uncertainties for Deep Learning. arXiv:1811.00908 [cs, stat]. arXiv: 1811.00908.

Thackeray, S. J., Henrys, P. A., Hemming, D., Bell, J. R., Botham, M. S., Burthe, S., Helaouet, P., Johns, D. G., Jones, I. D., Leech, D. I., Mackay, E. B., Massimino, D., Atkinson, S., Bacon, P. J., Brereton, T. M., Carvalho, L., Clutton-Brock, T. H., Duck, C., Edwards, M., Elliott, J. M., Hall, S. J. G., Harrington, R., Pearce-Higgins, J. W., Høye, T. T., Kruuk, L. E. B., Pemberton, J. M., Sparks, T. H., Thompson, P. M., White, I., Winfield, I. J., and Wanless, S. (2016). Phenological sensitivity to climate across taxa and trophic levels. Nature, 535(7611):241–245. Number: 7611 Publisher: Nature Publishing Group.

Thackeray, S. J., Sparks, T. H., Frederiksen, M., Burthe, S., Bacon, P. J., Bell, J. R., Botham, M. S., Brereton, T. M., Bright, P. W., Carvalho, L., Clutton-Brock, T., Dawson, A., Edwards, M., Elliott, J. M., Harrington, R., Johns, D., Jones, I. D., Jones, J. T., Leech, D. I., Roy, D. B., Scott, W. A., Smith, M., Smithers, R. J., Winfield, I. J., and Wanless, S. (2010). Trophic level asynchrony in rates of phenological change for marine, freshwater and terrestrial environments. Global Change Biology, 16(12):3304–3313. _eprint: https://onlinelibrary.wiley.com/doi/pdf/10.1111/j.1365-2486.2010.02165.x.

Tuia, D., Pozdnoukhov, A., Foresti, L., and Kanevski, M. (2012). Active Learning for Monitoring Network Optimization. In Spatio-Temporal Design, pages 285–318. John Wiley & Sons, Ltd.

UKCEH (2018). UKCEH Land Cover® plus Crops maps: UKCEH. © RSAC. © Crown Copyright 2007, Licence number 100017572.

Wu, W., May, R. J., Maier, H. R., and Dandy, G. C. (2013). A benchmarking approach for comparing data splitting methods for modeling water resources parameters using artificial neural networks. Water Resources Research, 49(11):7598–7614. _eprint: https://agupubs.onlinelibrary.wiley.com/doi/pdf/10.1002/2012WR012713.

Ye, L., Gao, L., Marcos-Martinez, R., Mallants, D., and Bryan, B. A. (2019). Projecting Australia’s forest cover dynamics and exploring influential factors using deep learning. Environmental Modelling & Software, 119:407–417.

Zheng, F., Maier, H. R., Wu, W., Dandy, G. C., Gupta, H. V., and Zhang, T. (2018). On Lack of Robustness in Hydrological Model Development Due to Absence of Guidelines for Selecting Calibration and Evaluation Data: Demonstration for Data-Driven Models. Water Resources Research, 54(2):1013–1030. _eprint: https://agupubs.onlinelibrary.wiley.com/doi/pdf/10.1002/2017WR021470.

Zhou, X.-L., Harrington, R., Woiwod, I. P., Perry, J. N., Bale, J. S., and Clark, S. J. (1995). Effects of temperature on aphid phenology. Global Change Biology, 1(4):303–313. Number: 4 Publisher: Wiley.

